# The condensin II/TOP-2 axis silences transcription during germline specification in *C. elegans*

**DOI:** 10.1101/2022.08.30.505898

**Authors:** Mezmur D. Belew, Emilie Chien, Matthew Wong, W. Matthew Michael

**Author notes:** denotes corresponding author. these authors contributed evenly to this work (Mezmur D. Belew, Emilie Chien).

## Abstract

In *C. elegans*, the germline is specified via a preformation mechanism that relies on the PIE-1 protein’s ability to globally silence mRNA transcription in germline precursor cells, also known as the P-lineage. Recent work from our group has identified additional genome silencing events in *C. elegans* during oogenesis and in starved L1 larvae, and these require the condensin II complex, topoisomerase II (TOP-2), and components of the H3K9me/heterochromatin pathway. Interestingly, silencing in oocytes also requires PIE-1, but this is not the case in starved L1s. Here, we ask if additional genome silencing components besides PIE-1 are required to repress gene expression in the P-lineage of early embryos, and we find that condensin II and TOP-2 are required and the H3K9me/heterochromatin pathway is not. We show that depletion of condensin II/TOP-2 activates the normally suppressed RNA polymerase II to inappropriately transcribe somatic genes in the P-lineage. We also present evidence that while both PIE-1 and condensin II/TOP-2 are required for genome silencing in the P-lineage, PIE-1 can silence transcription independently of condensin II/TOP-2 when misexpressed in somatic cells. Thus, in oocytes, all three genome silencing systems (TOP-2/condensin II, H3K9me, and PIE-1) are operational while in both early embryos and starved L1s two of the three are active (TOP- 2/condensin II and PIE-1 for early embryos, TOP-2/condensin II and H3K9me for starved L1s). Our data show that multiple, redundantly acting genome silencing mechanisms act in a mix and match manner to repress transcription at different developmental stages in the *C. elegans* germline.

## Introduction

Just like individual genes, the transcriptional output of entire genomes can be regulated in a signal-mediated manner. This is particularly true during germline development in *C. elegans,* where multiple cycles of whole genome activation and silencing occur (summarized in Fig 1). Starting with the gametes, we and others have shown that for both oocytes and spermatocytes transcription is globally silenced as these cells enter meiotic prophase (1–3). Silencing during gametogenesis requires three distinct pathways. One is comprised of topoisomerase II (TOP-2 in worms) and condensin II, which work together to compact chromatin and to silence gene expression (2,3). The second is comprised of components of the H3K9me/heterochromatin pathway, for example the methyltransferases SET-25 and MET-2, and is characterized by a large-scale buildup of H3K9me marks on chromatin as the genome is silenced (2). Lastly, the PIE-1 protein is also required for genome silencing during oogenesis (2). Upon fertilization, transcription continues to be repressed in the newly formed zygote, via the activities of the OMA- 1/2 proteins, and this continues through first mitosis into the 2-cell embryo (4). After the second mitosis, which forms the 4-cell embryo, OMA-1/2 are abruptly degraded and this allows transcriptional activation in the somatic cells ABa, ABp, and EMS. The germline precursor, P2, remains transcriptionally repressed due to the activity of PIE-1 (5,6). PIE-1 continues to repress transcription during germline specification in the so-called P lineage (P2, P3, and P4) until P4 divides to form the primordial germ cells Z2 and Z3. Upon division of P4 the PIE-1 protein is instantly degraded (5,7), and transcription is activated in the germline for the first time. Recent work for our group has shown that transcription continues in Z2/Z3 as the embryo hatches to form an L1 larva, but only if nutrients are present (8). If embryos hatch into a nutrient-free environment, then the energy sensor AMPK is activated and this promotes chromatin compaction and genome silencing in a manner dependent on the TOP-2/condensin II and H3K9me pathways (8). In this condition PIE-1 is dispensable for genome silencing as there is no detectable PIE-1 protein present in starved Z2/Z3 (E.C. and W.M.M., unpublished observation). Thus, for the germline, transcription is globally repressed during gametogenesis, as well as in 1- and 2-cell embryos. Transcription remains repressed in the germline progenitor P cells prior to activation in Z2/Z3, and can then be silenced again during L1 starvation and reactivated after feeding.

**Figure 1:**
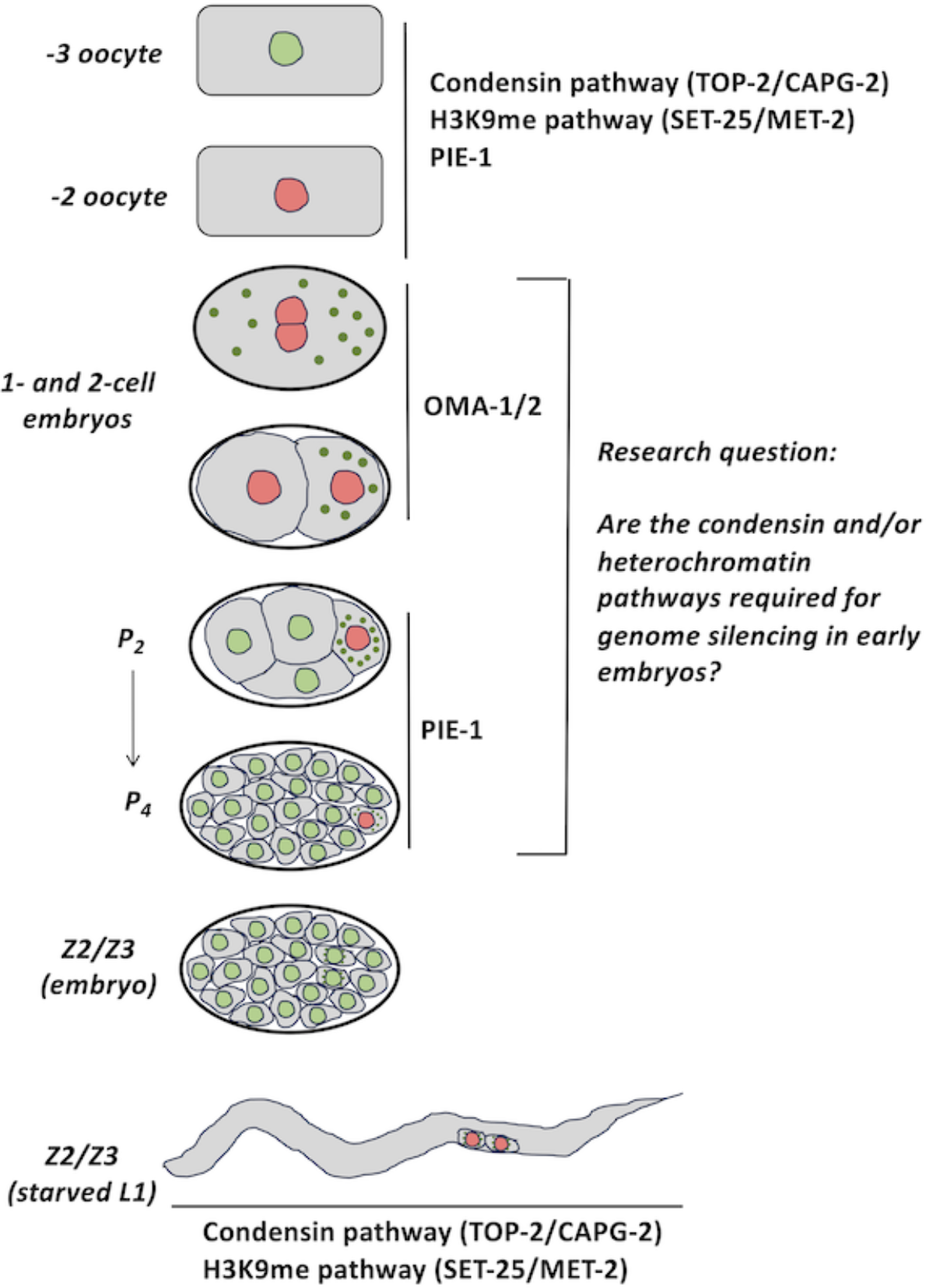
Schematic illustrating known genome silencing components during *C. elegans* germline development. Please see text for details.

The findings discussed above highlight the mechanistic plasticity in how germline transcription is controlled globally. Oocytes use TOP-2/condensin II, the H3K9me pathway, and PIE-1, and loss of any one of these pathways prompts unscheduled transcription. After fertilization, OMA-1/2 become essential for silencing in P0 and P1. The requirement for PIE-1 returns with the birth of P2, and this lasts until PIE-1 is destroyed upon division of P4. Later, in starved L1s, TOP-2/condensin II and the H3K9me heterochromatin pathway are required once again, and this time they act in a concerted manner. Thus, multiple systems are active in oocytes and Z2/Z3, however in the P-lineage, to date, we only know of single systems – OMA-1/2 for P0/P1 and PIE-1 for P2-P4. In this work we asked if the TOP-2/condensin II and/or the H3K9me heterochromatin pathways are required in the P-lineage for genome silencing, and found that the H3K9me heterochromatin pathway is dispensable while TOP-2/condensin II is required for P2-P4, but not for P0-P1. We also find that TOP-2/condensin II requires the germline progenitor state to silence the genome, and that PIE-1 does not. These findings increase our understanding of how whole-genome silencing occurs and highlight the importance of developmental context in regulating the ability of TOP-2/condensin II to perform this task.

## Materials and Methods

### C. elegans strains

N2 (wild-type), WMM2 *(ltls37 [(pAA64) pie-1p::mCherry::his-58 +unc-119(+)] IV; unc- 4(e120) top-2(it7ts) II)*, MT17463 *(set-25(n5021) III)*, WM330 *(pie-1(ne4301[pie-1::GFP]) III)*, AG275 *(top-2(av64)[TOP-2::3XFLAG] II)* and EG5175 *(pie-1(ne4301[pie-1::gfp]) III; mex-5(egx1[F294N & F339N]) IV)* strains were used in this study. Worms were maintained on 60-mm plates containing nematode growth media (NGM) seeded with the *E. coli* strain OP50 or HT115. Worms were grown at 20°C and propagated through egg preparation (bleaching) every 72 hours.

### Bacterial strains

OP50 bacteria served as the primary food source. It was grown in LB media containing 100 µg/ml streptomycin by shaking at 37°C overnight. 500 µl of the culture was seeded on Petri dishes containing NGM + streptomycin. HT115 bacteria grown in LB media containing 100 µg/ml carbenicillin and 12.5 µg/ml tetracycline and seeded on NGM + carbenicillin + tetracycline plates were also used as a source of food. Our RNAi strains were obtained from the Ahringer library and verified by Sanger sequencing. Bacteria containing dsRNA were streaked on LB-agar plates containing 100 µg/ml carbenicillin and 12.5 µg/ml tetracycline and incubated at 37°C overnight. Single colonies were then picked and grown in 25 ml LB cultures with 100 µg/ml carbenicillin and 12.5 µg/ml tetracycline. 500 µl of this culture was seeded on 60-mm Petri dishes containing 5 mM IPTG.

### Egg preparation

Bleach solution containing 3.675 ml H2O, 1.2 NaOCl, and 0.125 ml 10N NaOH was prepared. Adult worms were washed from plates with 5 ml of M9 minimal medium (22mM KH2PO4, 22mM Na2HPO4, 85mM NaCl, and 2mM MgSO4). Worms were centrifuged at 1.9 KRPM for 1 minute and the excess medium was removed, then the bleach solution was added. Eggs were extracted by vortexing for 30 seconds and shaking for 1 minute. This was done a total of 3 times and worms were vortexed one last time. Then the eggs were spun down at 1900 rpm for 1 minute and excess bleach solution was removed and the eggs were washed 3 times with M9 minimal medium.

### RNAi treatment

RNAi containing NGM plates were prepared as described in the “Bacterial strains” section. For double RNAi treatments, RNAi cultures were mixed at a 1:1 ratio by volume. HT115 cells transformed with an empty pL4440 vector was used as a negative control. RNAi conditions used in this study and tests for their efficacy is described below:

#### *top-2* RNAi

L1 worms were plated on HT115 food plates for the first 24 hours and were then moved to plates seeded with *top-2* RNAi for the remaining 48 hours. Embryonic lethality was observed at >90%.

#### *capg-2* RNAi

Worms were grown on HT115 food plates for the first 24 hours and were moved to plates containing *capg-2* RNAi for the remaining 48 hours. An embryonic lethality of 80%-100% was seen with this RNAi treatment.

#### *pie-1* RNAi

Worms were grown on plates containing *pie-1* RNAi for the entirety of their life cycle. An embryonic lethality of 100% was observed for this RNAi.

#### *mex-5/6* RNAi

Worms were grown on HT115 food plates for the first 48 hours of their life and were then switched to *mex-5/6* RNAi plates for the remaining 24 hours. An embryonic lethality of 90%-100% was observed for this RNAi treatment.

#### *mex-6*/control RNAi

Worms were grown on *mex-6*/control RNAi plates for the entirety of their life cycle. An embryonic lethality of 90%-100% was observed.

#### *mex-6/pie-1* RNAi

Worms were grown on *mex-6/pie-1* RNAi plates for the entirety of their life cycle. Am embryonic lethality of 90%-100% was observed.

### Antibodies and dilutions

#### RNAPIIpSer2

Rabbit antibody from Abcam (ab5095, Cambridge, Massachusetts) was used at a dilution of 1:100. <u>GFP</u>: mouse Mab #3580, from EMD Millipore, was used at 1:500. <u>FLAG</u>: Mouse antibody F1804 from EMD Millipore was used at 1:1000. <u>H3K9me3</u>: Rabbit antibody from Abcam (ab176916, Cambridge, Massachusetts) was used at a dilution of 1:1000. <u>P-granules</u>: mouse Mab OIC1D4 (Fig 1A) or mouse Mab K76 (remainder of study), both from the Developmental Studies Hybridoma Bank, were used at a dilution of 1:10 or 1:20, respectively. <u>Secondary antibodies</u>: Alexa Fluor conjugated secondary antibodies from Invitrogen (Thermofisher Scientific, Waltham, Massachusetts) were used at a dilution of 1:200.

### Immunofluorescence staining

Adult worms were first washed off plates with 10 ml of M9 minimal medium and washed 3 more times. Then, they were centrifuged at 1.9 KRPM and the excess medium was removed. 20 µl of media containing about 50 worms were spotted on a coverslip and 3 µl of anesthetic (20mM Sodium Azide and 0.8M Tetramisole hydrochloride) was added to immobilize them. Worms were dissected using 25Gx5/8 needles (Sigma Aldrich, St. Louis, Missouri). To release early embryos, worms were cut once midway through their length. To release gonads for oocyte staining, worms were cut twice, once near the head and once near the tail. The coverslip was then mounted onto poly-L-lysine covered slides and let rest for 5 minutes. Slides were put on dry ice for 30 minutes. Samples were then freeze-cracked by flicking the coverslips off for permeabilization.

For RNAPIIpSer2 staining experiments, once samples are permeabilized, slides were put in cold 100% methanol for 2 minutes and then fixing solution (0.08M HEPES pH 6.9, 1.6mM MgSO4, 0.8mM EGTA, 3.7% formaldehyde, 1X phosphate-buffered saline) for another 30 minutes. After fixing, slides were washed three times with TBS-T (TBS with 0.1% Tween-20) and were blocked for 30 minutes with TNB (containing 100mM Tris-HCl, 200mM NaCl, and 1% BSA). Primary antibodies were then applied at the dilutions described above in TNB and slides were incubated at 4°C overnight.

For H3K9me3 and GFP staining experiments, permeabilized samples were put in cold 100% methanol for 10 seconds and then fixing solution (0.08M HEPES pH 6.9, 1.6mM MgSO4, 0.8mM EGTA, 3.7% formaldehyde, 1X phosphate-buffered saline) for 10 minutes. After fixing, slides were washed three times with TBS-T (TBS with 0.1% Tween-20) and were blocked for 2 hours with TNB (containing 100mM Tris-HCl, 200 mM NaCl, and 1% BSA) supplemented with 10% goat serum. Primary antibodies were then applied at the dilutions described above in TNB and slides were incubated at 4°C overnight. On the next day, the slides were washed 3 times with TBS and slides were incubated with secondary antibodies and Hoechst 33342 dye for 2 hours at room temperature. Slides were washed 3 times with TBS, mounting medium (50 % glycerol in PBS), and coverslips were applied and sealed with Cytoseal XYL (Thermofisher).

### HCR (In Situ Hybridization Chain Reaction)

A kit containing a DNA probe set, DNA hybridized chain reaction (HCR) amplifier hairpins, and hybridization, wash, and amplification buffers were purchased from Molecular Instruments (molecularinstruments.com). Genes that were examined were *vet-6* and *F58E6.6*. DNA was visualized with Hoechst 33342 dye. Embryos were prepared by bleaching and were immediately spotted on poly-L-lysine coated slides. Coverslips were applied and slides were freeze-cracked to permeabilize samples. Immediately after freeze-cracking, 500 µl of 100% ice-cold methanol was applied over the samples. Slides were dried off by tilting slides, 1 ml of 4% paraformaldehyde (PFA) was added and samples were incubated in a humidity chamber for 10 minutes. Samples were then washed 3 times with 100 µl of PBS-T. A 1:1 solution of probe hybridization buffer (PHB) and PBS-T was added to the samples, and they were incubated for 5 minutes at room temperature. Samples were then prehybridized with PHB for 30 minutes at 37°C and DNA probes (at a final concentration of 2 picomoles per 500 µl of PHB) were added to the samples and were incubated overnight at 37°C. The next day, samples were washed 4 times with probe wash buffer (PWB) at 37°C with 15 minutes of incubation for each wash. They were then washed three more times with 5xSSCT at room temperature. Samples were pre-amplified with Amplification buffer for 30 minutes at room temperature. Probe amplifier hairpins were snap cooled by heating to 95°C for 90 seconds and putting in a dark drawer for 30 mins then were added to the sample. Worms were incubated with the hairpins overnight in a dark drawer. On the third day, samples were washed with 5xSSCT and incubated with Hoechst-33342 (1:5000 dilution) for 15 minutes. Finally, samples were mounted on poly-L-lysine coated slides and imaged.

### Immunofluorescent imaging

All slides were imaged using an Olympus Fluoview FV1000 confocal microscope using Fluoview Viewer software. A magnification of 600x (60x objective and 10x eyepiece magnifications) was used. Laser intensity was controlled for experiments to achieve consistency among samples.

## Results

### One- and two-cell embryos do not require either SET-25 or TOP-2 for genome silencing

We first examined one- and two-cell embryos, which previous work has shown to be reliant on OMA-1/2 for transcriptional repression (4). One hallmark of silencing in both oocytes and starved Z2/Z3 is a large accumulation of H3K9me3 marks on chromatin, as inferred by immunofluorescence microscopy (2,8). In *C. elegans*, oocytes are named by their position relative to the spermatheca in the tube-shaped gonad, where -1 is closest. Starting at the -5 position, H3K9me3 marks begin to accumulate on chromatin such that at the -2 and -1 positions H3K9me3 signals overlap with much of the DAPI-based DNA signal (ref 2, see Fig 2A). By contrast, in zygotes, the maternal pronucleus (as well as its paternal counterpart) showed greatly reduced H3K9me3 signal intensity, manifesting as small patches on otherwise barren chromatin (Fig 2B). This was true of both newly fertilized zygotes and those that were nearing mitosis (Fig 2B). Interestingly, the polar bodies retained strong H3K9me3 signals (Fig 2B). These data suggest that, upon fertilization, H3K9me3 marks are rapidly erased from maternal chromatin, but this does not occur on polar bodies as they have been ejected from the zygote’s cytoplasm and thereby escape erasure.

**Figure 2:**
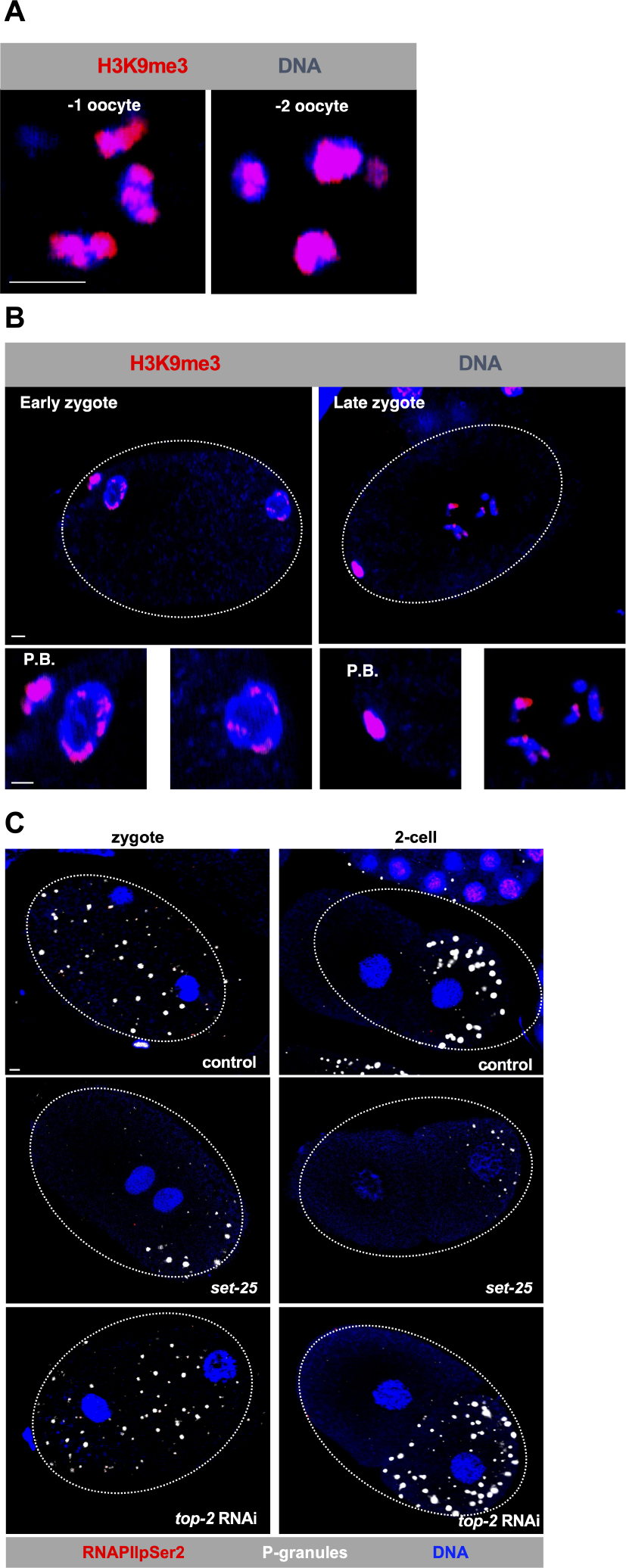
No role for either the H3K9me heterochromatin pathway or the TOP-2/condensin II pathway in transcriptional repression in zygotes or two-cell embryos. (A) Wild-type oocytes were fixed and stained for H3K9me3 (red) and DNA (blue). Shown are magnified views of the -1 and -2 oocytes, which are the oocytes most proximal and second-most proximal to the spermatheca, respectively. The H3K9me3 signal overlaps with the Hoechst signal extensively. Scale bar represents a length of 2 µm. (B) The indicated zygotes from N2 worms were fixed and stained for H3K9me3 (red) and DNA (blue). Shown below the entire embryos are magnified views of the pronuclei and polar bodies (PB). Note that the H3K9me3 signal overlaps with the Hoechst signal more extensively on the polar bodies than within the pronuclei. Scale bar represents a length of 2 µm. (C) Zygotes and two-cell embryos were fixed and stained for RNAPIIpSer2 (red), DNA (blue), and P-granules (white). No active RNAPII was observed in the more than 20 samples we examined for both one- and two-cell embryos over 2 replicates. Note the RNAPIIpSer2 signals in the adjacent, older embryo in the upper right panel – this shows that the staining worked efficiently. Scale bar represents a length of 2 µm.

Our previous work has shown that the SET-25 methyltransferase is responsible for the accumulation of H3K9me3 marks on both oocyte and starved Z2/Z3 chromatin, and this enzyme is also required for transcriptional repression in these cells (2,8). To ask if SET-25 plays a role in transcriptional silencing in one- or two-cell embryos we monitored *set-25^n5021^* mutant embryos for active RNA polymerase II (RNAPII). For this we used immunofluorescence staining with an antibody that detects a phospho-serine 2 epitope on the RNAPII carboxyl-terminal domain (CTD). The presence of this RNAPIIpSer2 signal in nuclei is a marker for actively elongating RNAPII (9), and we and others have used this approach extensively to identify transcriptionally active nuclei within a variety of *C. elegans* tissues (1–4, 10–16). The antibody that we are using, a rabbit polyclonal that recognizes RNAPIIpSer2, has previously been validated by us for use in *C. elegans* embryos (8). The samples were also stained with an antibody recognizing P-granules, as these structures are specific to P-cells and can therefore be used to distinguish them from somatic cells. As shown in Fig 2C, *set-25* mutants were still fully capable of repressing transcription in both one- and two-cell embryos. Next, we used RNAi to deplete TOP-2. Our previous work, with these same RNAi conditions, has shown that loss of TOP-2 promotes unscheduled transcription in oocytes, spermatocytes, and starved Z2/Z3 (2,3,8). However, in one- or two-cell embryos, TOP-2 depletion had no impact on transcriptional repression (Fig 2C). We conclude that the H3K9me pathway is not required for transcriptional silencing in one- or two-cell embryos, and that the same is true for TOP-2.

### Loss of TOP-2 and condensin II function results in active RNA polymerase II (RNAPII) in the P-lineage of early embryos

We next examined P2 and P3 for the presence of H3K9me3 as well as a requirement for SET-25 in transcriptional repression, with the idea that the H3K9me pathway might become required for silencing upon loss of OMA-1/2. This was not the case, however, as there were no detectable differences in the amount of H3K9me3 marks on P2 or P3 relative to their somatic sister cells (EMS and C, respectively, see Fig S1A). In addition, transcription was not derepressed in P2 in *set-25^n5021^* mutant embryos (Fig S1B). We thus turned our attention to the possibility that the TOP-2/condensin II pathway might play a role in genome silencing in P2, P3, and P4. For this we again stained embryos for RNAPIIpSer2 and P-granules, as we did in Fig 1C. In control 4-cell embryos the expected pattern emerged – signal could be seen in the somatic ABa, ABp, and EMS cells, but not P2 (Fig 3A). We next depleted either TOP-2 or the CAPG-2 subunit of condensin II. Interestingly, in samples exposed to either *top-2* or *capg-2* RNAi, we could now detect RNAPIIpSer2 in P2, and the same was true of samples exposed to *pie-1* RNAi (Fig 3A). We also examined the remainder of the P-lineage (P3 and P4) and in both cases we could detect active transcription after depletion of either TOP-2, CAPG-2, or PIE-1 (Fig 3A). Based on these data, we conclude that TOP-2/condensin II is dispensable for transcriptional repression in P0 and P1, but for P2, P3, and P4 the pathway is required to silence gene expression. We note that in Fig 3A the P-granules would occasionally manifest as a yellow color, instead of green as would be expected by the combination of primary and secondary antibodies used to detect them. We believe this is due to occasional cross-reactivity between the secondary antibody used to detect RNAPIIpSer2 (Alexa 555 goat anti-rabbit, ThermoFisher #A21428) and the primary antibody used to detect P-granules (Mab OIC1D4, from ATCC). This was not an issue when we used Mab K76 to detect P-granules (see Figs 2C and S1B) and because P-granules are localized outside of the nucleus the occasional cross-reactivity between Alexa 555 goat anti-rabbit and Mab OIC1D4 did not impact data collection or interpretation.

**Figure 3:**
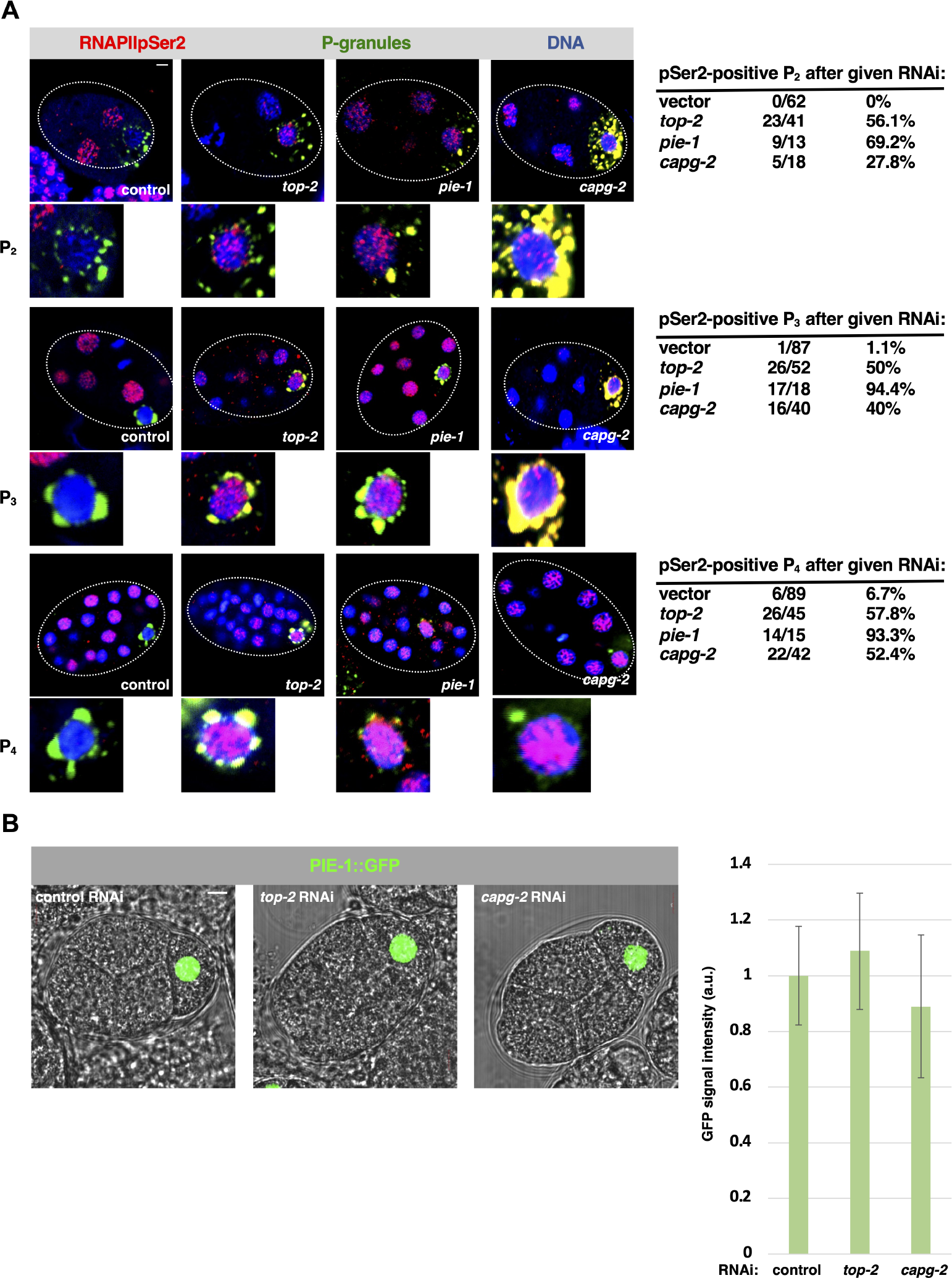
Loss of TOP-2 or CAPG-2 allows for active RNAPII in the P-lineage of early embryos. (A) Early embryos from N2 worms treated with control, *top-2*, *capg-2,* or *pie-1* RNAi were fixed and stained for RNAPIIpSer2 (red), DNA (blue) and P-granules (green). Shown on top are the entire embryos and the P-cell is magnified below. Quantification of the data is shown at right. We note that the P-granules in this data set would occasionally manifest as a yellow signal, as opposed to the green signal expected, from the combination of primary and secondary antibodies used to detect them. Please see Results for an explanation for this. (B) Fluorescence and phase-contrast microscopy was used to image four-cell embryos expressing PIE-1::GFP after treatment with the indicated RNAi. The graph shows GFP signal quantification across the three samples. For signal quantification, confocal slices corresponding to maximal signal intensity were analyzed using ImageJ software to measure pixel density. GFP signal intensity was normalized to control. Scale bar represents a length of 5 µm.

It was important to check if *top-2* or *capg-2* depletion reduced PIE-1 expression in P2-P4, as this would explain the appearance of RNAPIIpSer2 in these cells. To do so we used a strain harboring PIE-1 tagged with GFP at the endogenous locus (referred to as PIE-1::GFP, refs 17,18). PIE-1::GFP is expressed normally and retains PIE-1 function (17,18). We measured GFP signal intensity in P-cells after control, *top-2*, or *capg-2* RNAi and, as shown in Fig 3B, PIE-1::GFP expression in P2 was not noticeably altered after depletion of TOP-2 or CAPG-2. Thus, the mechanism by which TOP-2/condensin II silences gene expression in the P-lineage does not involve the control of PIE-1 expression levels.

### Aberrant expression of EMS-specific genes is observed in the P-lineage when TOP- 2 and condensin II are depleted

Data presented thus far demonstrate aberrant RNAPII activity in the P-lineage after depletion of TOP-2 or CAPG-2 (Fig 3A). To obtain additional evidence for the misregulation of gene expression in these cells, we performed the *in-situ* hybridization chain reaction (HCR), also known as RNA-FISH (19), to detect and localize mRNAs in early embryos. We identified a transcript, F58E6.6, that is found predominantly in EMS under normal conditions, with only 10- 20% of the wild-type samples showing transcript signals in ABa, ABp, or P2 (Fig 4A, quantification is shown to the right in Fig 4B on a per embryo basis). After TOP-2 or CAPG-2 depletion, however, 50% and 82%, respectively, of the embryos expressed F58E6.6 in P2 (Fig 4A&B). As a positive control, we depleted PIE-1 and found that F58E6.6 was expressed in P2 in 5 out of 5 samples (Fig 4A&B). These data show that loss of either TOP-2/condensin II or PIE-1 allows unscheduled transcription of F58E6.6 in P2. We also examined another EMS-specific transcript, the previously reported *vet-6* (6). We observed that *vet-6* was also misexpressed in P2 upon depletion of TOP- 2, CAPG-2, or PIE-1 (Fig S2), however the effect of CAPG-2 depletion was not as pronounced as that seen for the F58E6.6 transcripts in Fig 4. Based on these data, we conclude that depletion of TOP-2 and condensin II results in the misexpression of EMS genes in the P-lineage of early embryos.

**Figure 4:**
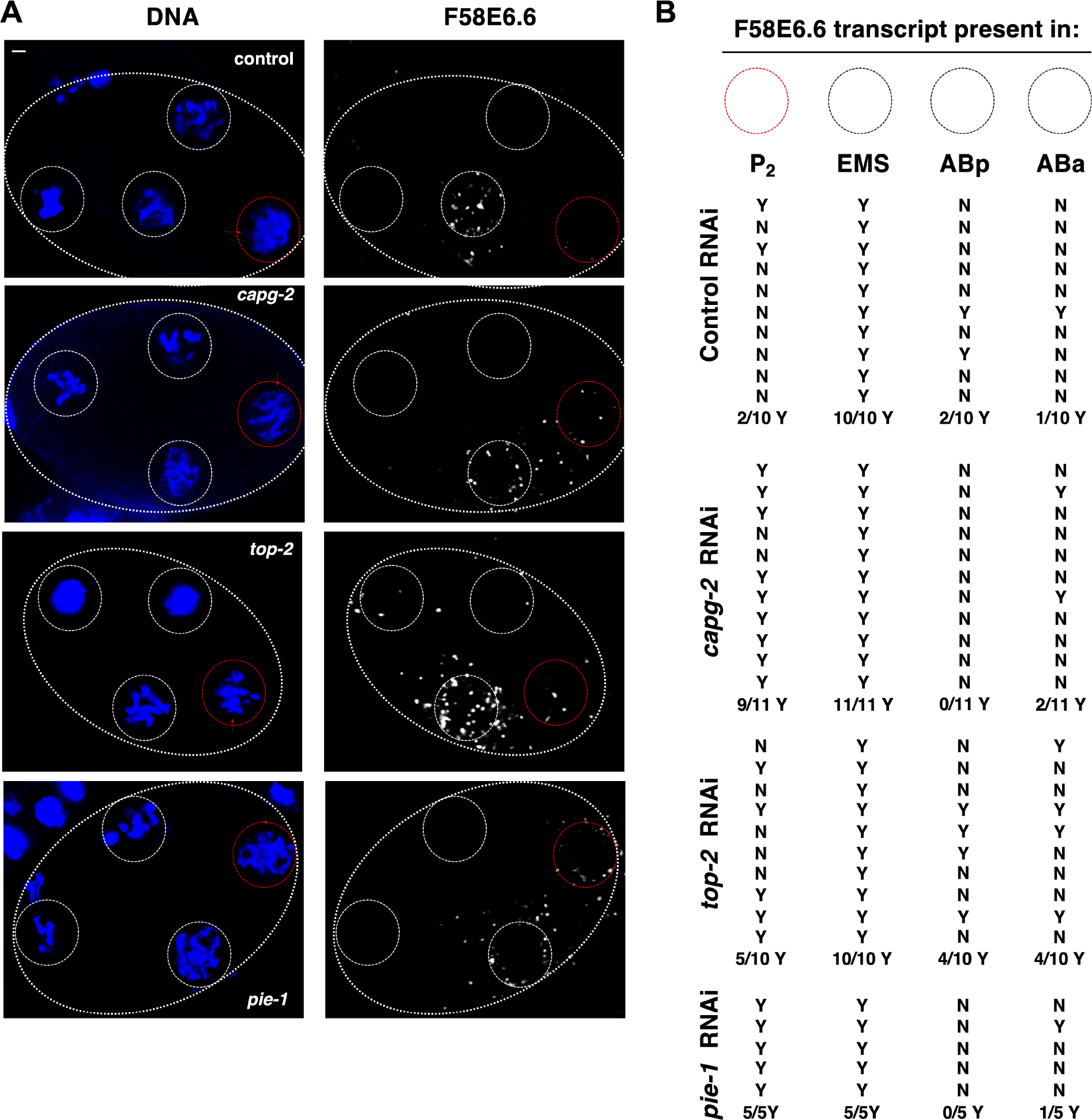
EMS-specific gene is aberrantly expressed in the P2 cell of 4-cell embryos after loss of TOP-2 or CAPG-2. (A) Four-cell embryos from N2 animals treated with control, *capg-2*, *top-2,* or *pie-1* RNAi were fixed and HCR was performed to probe for *F58E6.6* mRNA (white). DNA was stained using Hoechst-33342 (blue). Representative images are shown. White dashed circles represent the EMS cell and the red dashed circles represent P2. (B) Percentage of samples with *F58E6.6* transcripts in EMS P2 (n=40).

### PIE-1 can repress transcription outside of the P-lineage whereas TOP-2 cannot

Our work has shown that genome silencing in the P2, P3, and P4 cells requires both PIE-1 and the TOP-2/condensin II pathway. PIE-1 is expressed exclusively in the P-lineage, however TOP-2 and condensin II are likely expressed in all blastomeres of the early embryo. To confirm this for TOP-2 we used a strain where TOP-2 contains a FLAG tag at the endogenous locus (referred to as TOP-2::FLAG). TOP-2::FLAG retains TOP-2 function (20) and is indeed present in all 4 cells of the 4-cell embryo (Fig S3A). This raises the question of why TOP-2 can repress transcription in P-cells but not somatic cells. One possibility is that TOP-2 requires the presence of PIE-1 to silence the genome. We have previously shown that this is true in oocytes, where loss of either PIE-1 or TOP-2 prevents transcriptional repression (2). By contrast, PIE-1 is not expressed in the Z2/Z3 PGCs and TOP-2 is nonetheless required for genome silencing during L1 starvation (8). These findings show that TOP-2 can repress transcription through both PIE-1 dependent and independent mechanisms.

To discern the relationship between PIE-1 and TOP-2 in early embryos we sought a means of misexpressing PIE-1 in the early embryo. Previous work has shown that depletion of the translational regulators MEX-5 and MEX-6 allows PIE-1 expression in all cells of the 4-cell embryo (21). The loss of PIE-1 asymmetry in *mex-5/6* RNAi embryos is accompanied by loss of somatic asymmetries as well, e.g. the SKN-1 protein that is normally expressed only in the somatic EMS cell is also found in all four cells after *mex-5/6* RNAi (21). Thus, depletion of MEX-5/6 results in blastomeres assuming a merged somatic and germline precursor state. We confirmed the effect of *mex-5/*6 RNAi on PIE-1 localization by using the strain expressing PIE-1::GFP. As shown in Fig S3B, in control samples PIE-1::GFP is expressed solely in P2, whereas loss of MEX-5/6 causes a redistribution to all four blastomeres, as expected. Furthermore, when samples were stained for RNAPIIpSer2, control embryos displayed signals in EMS but not P2, as expected, while the *mex- 5/*6 RNAi samples did not show active RNAPII in any cells, consistent with the misexpression of PIE-1::GFP. Thus, loss of MEX-5/6 causes misexpression of PIE-1::GFP in all blastomeres, and also represses transcription in all blastomeres. To determine if misexpressed PIE-1::GFP is the reason that *mex-5/6* embryos are blocked for transcription we used a strain expressing both an RNA-binding deficient form of MEX-5 (*mex-5^egx1^*) and PIE-1::GFP (18). This strain is disabled for MEX-5 function but remains viable through MEX-6 activity (18). We treated this strain with *mex-6* RNAi together with either control or *pie-1* RNAi to create conditions where animals were depleted of either MEX-5/6 activity alone or MEX-5/6 <u>and</u> PIE-1::GFP activity. Transcriptional activity in four-cell embryos was then assessed via RNAPIIpSer2 staining. As shown in Fig S3C, in samples treated with *mex-6* plus control RNAi, PIE-1::GFP was expressed in all four blastomeres and RNAPIIpSer2 was attenuated. By contrast, in samples exposed to RNAi against both MEX-5 and GFP-PIE-1, GFP signals were reduced and RNAPIIpSer2signals were easily detected in all four blastomeres. We conclude that the pan-embryo transcriptional repression observed after loss of MEX-5/6 activity is due to misexpression of PIE-1::GFP.

To examine a role for TOP-2 in promoting transcriptional repression by misexpressed PIE- 1, we used a temperature-sensitive allele of *top-2* (20) in combination with *mex-5/6* RNAi and examined RNAIIpSer2 in 4-cell embryos at both the permissive temperature (15°C) and the non-permissive temperature (24°C). For control RNAi, we observed that inactivation of TOP-2 caused the appearance of RNAIIpSer2 in P2, as expected, whereas samples raised at the permissive temperature were devoid of RNAIIpSer2 in P2 (Fig 5A upper panels). Interestingly, after *mex-5/6* RNAi, transcription was attenuated in all four blastomeres, regardless of the activity state of TOP- 2 (Fig 5A, bottom panels). Taken together, our data show that inactivation of PIE-1 in a MEX-5/6 deficient condition prevents pan-embryo transcriptional repression (Fig S3C), whereas inactivation of TOP-2 in MEX-5/6 deficient samples does not (Fig 5A). These findings, in turn, suggest that TOP-2 requires the germline progenitor state for genome silencing in early embryos and that when this state is perturbed then TOP-2 loses its ability to repress transcription (summarized in Fig 5B).

**Figure 5:**
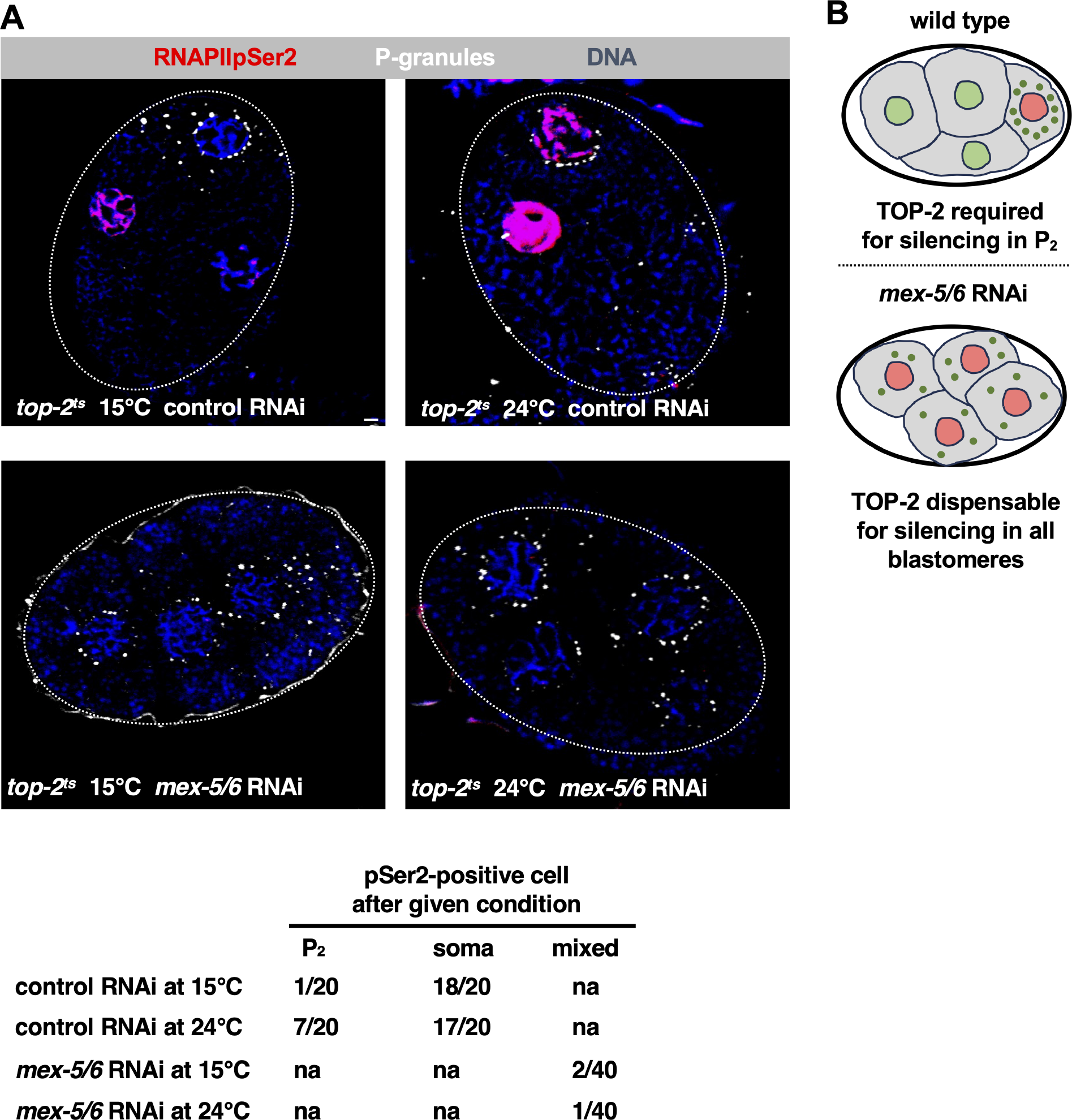
PIE-1 does not require TOP-2 to repress transcription in *C. elegans* early embryos. (A) Four-cell embryos from WMM2 (*top-2^ts^*) animals treated with either control or *mex-5/6* RNAi were optionally shifted to the nonpermissive temperature of 24°C for 24 hours or left at the permissive 15°C. Then samples were fixed and stained for RNAPIIpSer2 (red), DNA (blue), and P-granules (white). 20 samples were analyzed over 2 independent replicates. Attenuation of TOP-2 via temperature shift does not reverse the transcriptional silencing by PIE-1 in *mex-5/6* RNAi embryos. Quantification of the data is presented below, where “soma” indicates the EMS cell in control RNAi samples. Scale bar represents a length of 2 µm. (B) Cartoon of TOP-2 requirement for transcriptional silencing at the 4-cell embryonic stage. Nuclei colored in red indicate transcriptionally silenced cells.

## Discussion

In this study we focused on the early embryonic P-lineage with the goal of determining if the TOP-2/condensin II and/or H3K9me/heterochromatin genome silencing pathways are required for transcriptional repression in these cells. Previous work has revealed that two different genome silencers, OMA-1/2 and PIE-1, are active in the P-lineage, with OMA-1/2 responsible for P0 and P1 (as well as AB) and PIE-1 taking over for P2-P4 (Fig 1). Our work has added to this profile with the demonstration that TOP-2 and condensin II are needed for efficient silencing in P2-P4, but not P0 or P1, and that the H3K9me/heterochromatin pathway is not required at all in the P-lineage. We base these conclusions on the following data. For TOP- 2/condensin II we show that depletion of either TOP-2 or the condensin II subunit CAPG-2 results in the appearance of RNAPIIpSer2 signals in P2, P3, and P4 (Fig 3A) and inappropriate transcription of the somatic F58E6.6 and *vet-6* genes in P2 (Figs 4 and S2). We note that the impact of TOP-2 or CAPG-2 depletion on transcription in the P-lineage was not as profound as depletion of PIE-1 (Figs 3,4, and S2), and this is likely due to the fact that we use mild RNAi conditions for TOP- 2/CAPG-2, relative to PIE-1, as stronger conditions for TOP-2/CAPG-2 result in torn, cut, and malformed nuclei, as has been reported previously (22–24). Although we tested just one condensin II subunit in this work, our previous work had shown that depletion of KLE-2, in addition to CAPG-2, disrupts silencing in starved Z2/Z3 (8), and thus it seems likely that the *capg- 2* phenotypes reported here represent a loss of condensin II function and not a condensin II independent function of CAPG-2.

For the H3K9me/heterochromatin pathway we focused on the SET-25 methyltransferase, which we have previously shown to be required for genome silencing in both oocytes and in starved Z2/Z3 (2,8). We could not reveal a role for SET-25 in P-lineage silencing (Figs 2 and S1). Furthermore, unlike oocytes and starved Z2/Z3, we did not observe a hyper-accumulation of H3K9me3 marks in P-lineage nuclei (Figs 2 and S1), adding further support to the conclusion that the H3K9me/heterochromatin pathway is dispensable for genome silencing in early embryos.

Our work, presented here and elsewhere, has revealed a role for the TOP-2/condensin II pathway in transcriptional repression in three different contexts – oocytes and spermatocytes (refs 2-3 and Fig 6A ), starved Z2/Z3 PGCs (ref 8 and Fig 6B) and in the P2, P3, and P4 cells of early embryos (this work and Fig 6C). Mechanistically, available data suggest that TOP-2/condensin II silences transcription in oocytes via chromatin compaction, which could block access of the transcriptional machinery to gene promoters (“promoter occlusion” in Fig 6A). The same mechanism also applies to starved Z2/Z3, as we have shown that starvation triggers extreme chromatin compaction and that when compaction fails then nuclei remain transcriptionally active (ref 8 and Fig 6B). How the TOP-2/condensin II pathway silences transcription in P2-P4 is not clear as the requirement for DNA replication during cell division seemingly precludes the establishment of a stably compacted chromatin state. Another important question raised by our findings is why, in the early embryo, TOP-2/condensin II is only active for transcriptional repression in P-cells, despite being ubiquitously expressed. Indeed, we have shown here that when P-cell identity is lost, then PIE-1 no longer requires the presence of TOP-2 to block transcription, as it does in both the P-lineage and in oocytes (this work and ref 2). Thus, it appears that the there is something about the germline or germline progenitor state that transforms TOP- 2/condensin II into a genome silencing system, and further work is needed to understand how TOP-2/condensin II is differentially regulated in germline versus soma.

**Figure 6:**
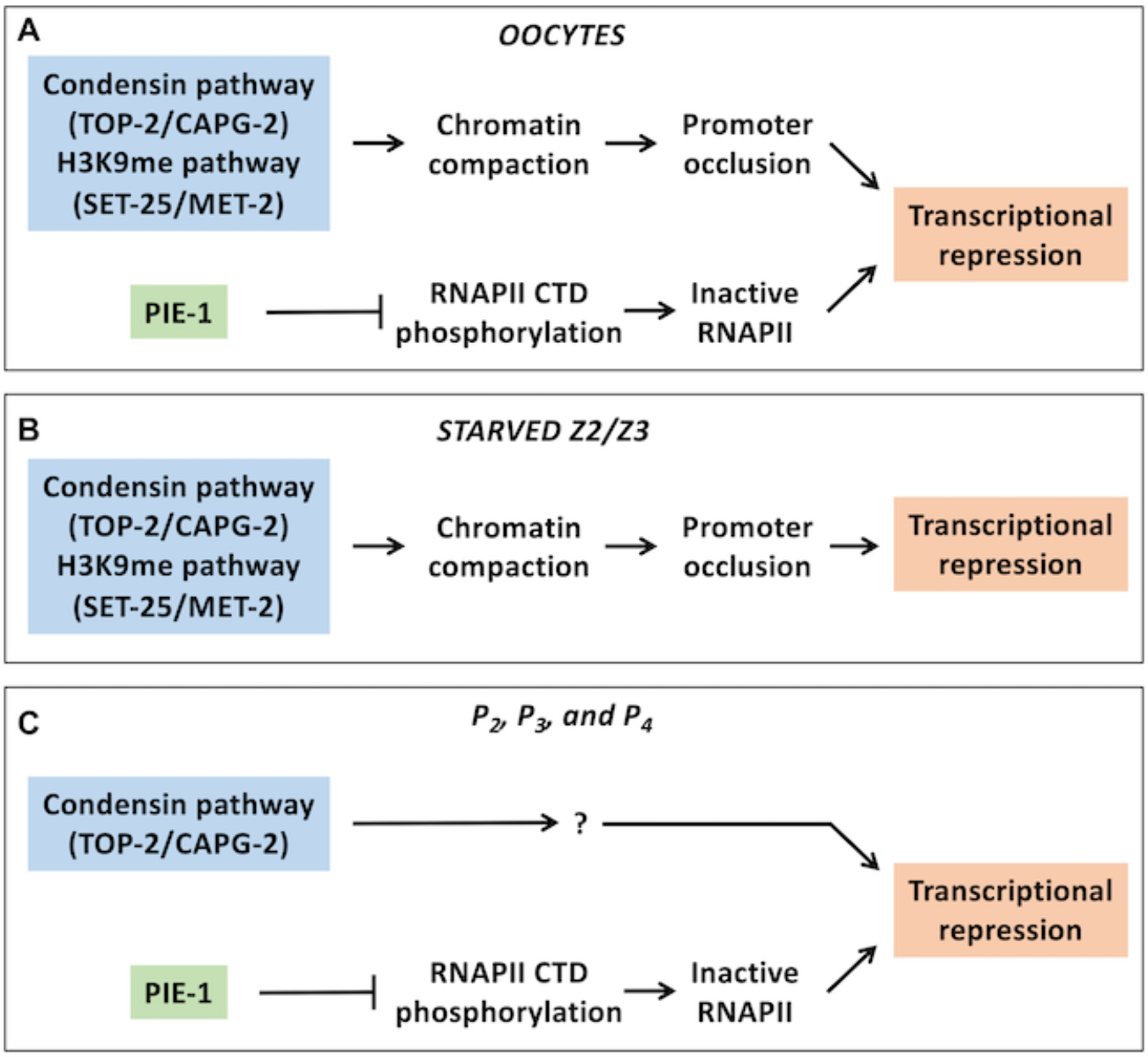
Summary of genome silencing components at different developmental stages in the germline. (A) In oocytes, TOP-2/CAPG-2 and the H3K9me pathway repress transcription via chromatin compaction while PIE-1 may work through control of RNAPII CTD phosphorylation. (B) In starved L1 larvae, TOP-2/CAPG-2 and the H3K9me pathway are required for transcriptional repression of Z2/Z3. (C) In P-lineage cells, TOP-2/CAPG-2 works independently of PIE-1 to promote transcriptional repression, although the precise mechanism by which TOP-2/CAPG-2 works is not yet known.

Lastly, our work has also revealed an intriguing connection between PIE-1 and the TOP- 2/condensin II pathway during germline genome silencing. We found that loss of either pathway prevents silencing in both oocytes and in P2-P4, suggesting that the two systems are dependent on one another as opposed to acting redundantly. The mechanism by which PIE-1 blocks transcription is unclear. Early work suggested that PIE-1 competes with the RNAPII CTD for binding to the cyclin T component of the CDK9-cyclin T complex that is required for phosphorylation and activation of RNAPII (25,26). PIE-1 has been proposed to titrate cyclin T away from RNAPII, thereby suppressing the CTD phosphorylations that are needed to trigger RNAPII elongation. More recent work, however, has shown that PIE-1 in the adult germline controls SUMOylation of chromatin remodeling factors, such as the NuRD complex component and the histone deacetylase HDA-1 (27). Thus it may be that PIE-1 connects to the TOP- 2/condensin II pathway through chromatin as opposed to direct regulation of RNAPII.

## Acknowledgements

This work was funded by NIH R01 award (R01GM127477) to W.M.M. We are grateful to Craig Mello and Erik Griffith for their kind gifts of worm strains.

## Supporting information figures and legends

**Figure S1:**
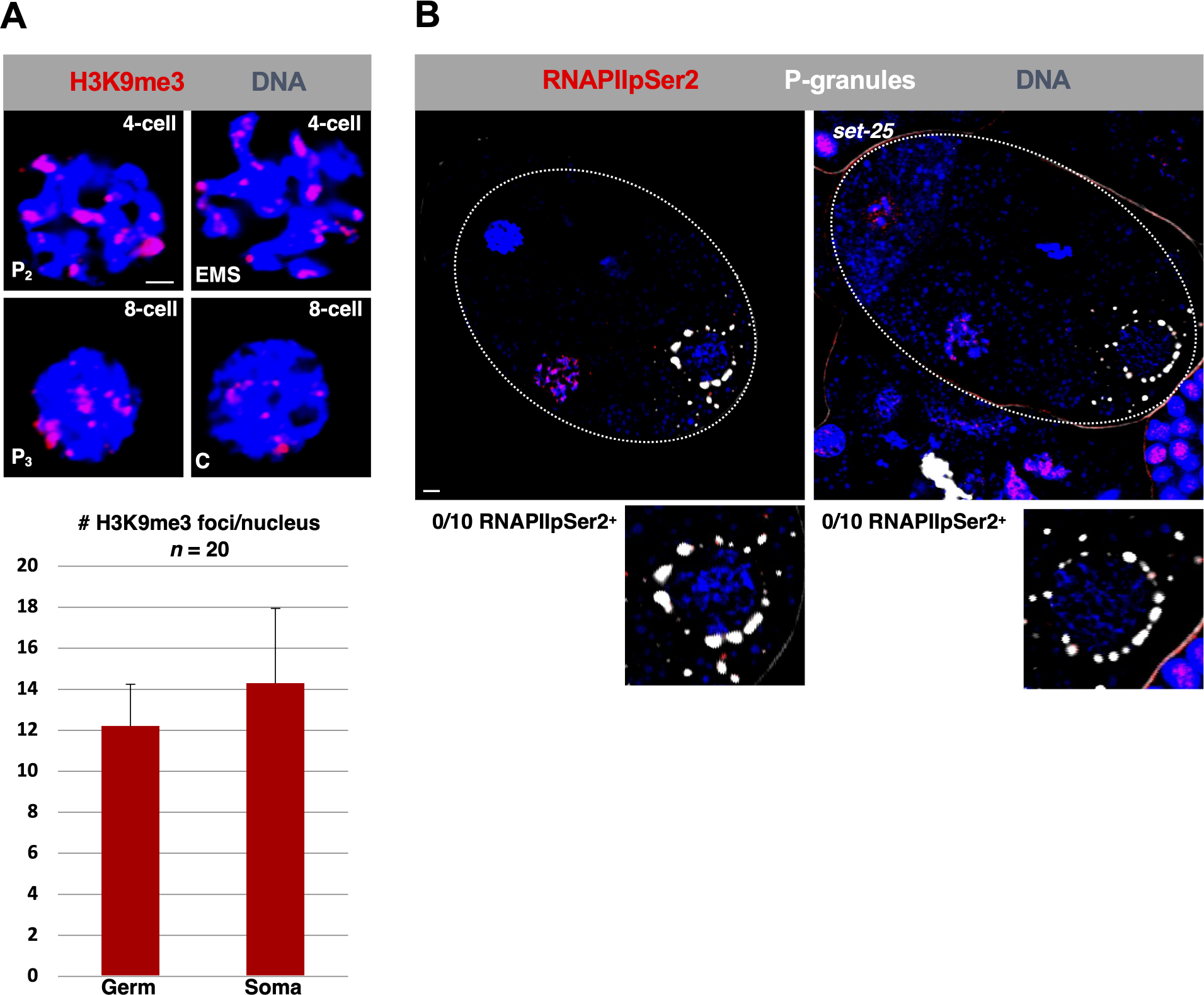
RNAPII-mediated transcription is still repressed in P2 after loss of SET-25. (A) Four- and eight-cell embryos were examined were fixed and stained for H3K9me3 (red) and DNA (blue). Quantification of H3K9me3 foci for P2 and P3 (Germ) or EMS and C (Soma) is shown below. Scale bar represents a length of 2 µm. (B) Four-cell embryos, either wild type (N2) or *set-25* mutants (strain MT17463), were fixed and stained for RNAPIIpSer2 (red) and P-granules (blue). 10 samples were analyzed. Both wild-type and *set-25* mutants lacked RNAPIIpSer2 signal in P2. Scale bar represents a length of 2 µm.

**Figure S2:**
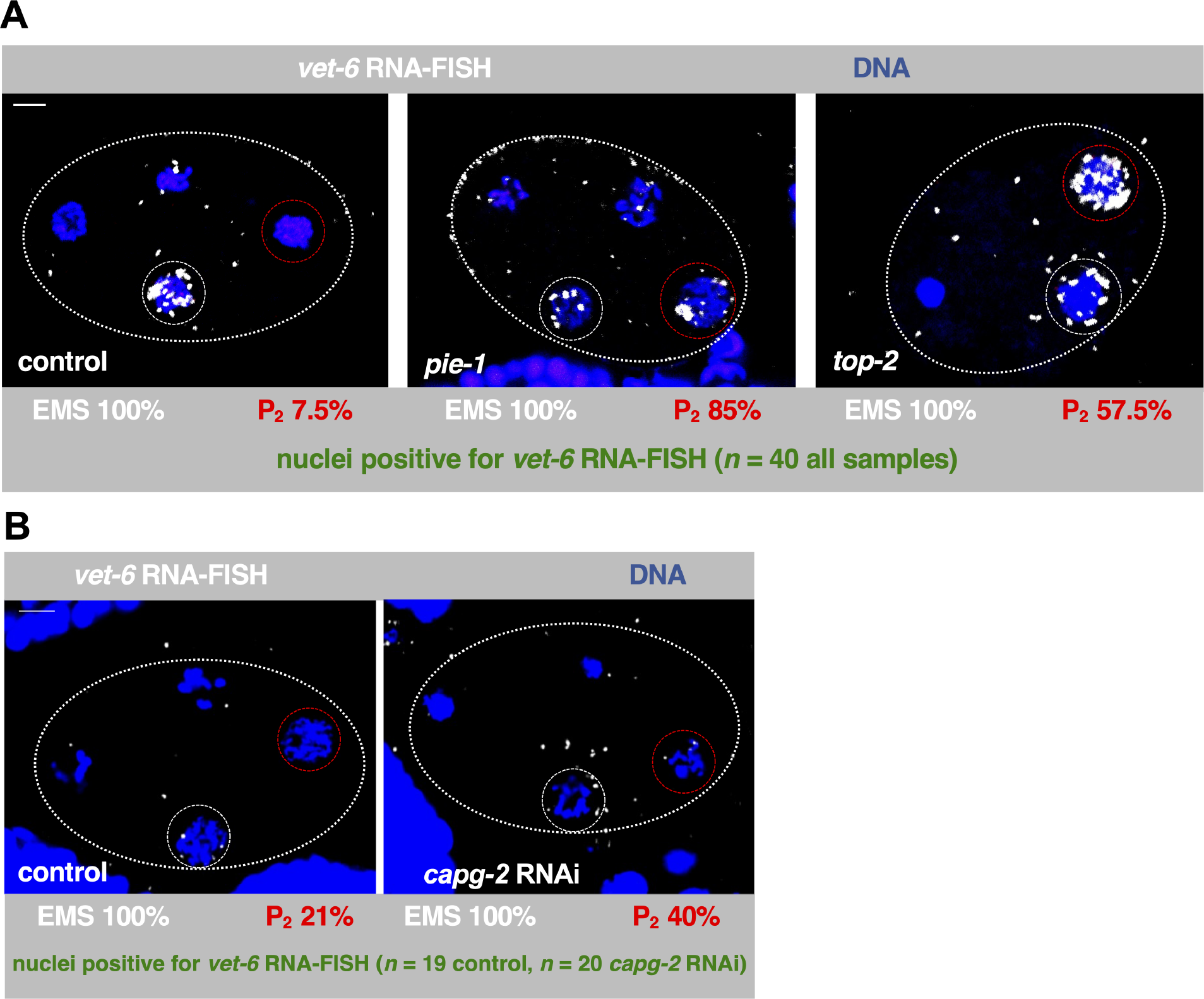
EMS specific gene is aberrantly expressed in the P2 cell of 4-cell embryos after loss of TOP-2 but not CAPG-2. (A) HCR was performed using 4-cell embryos from N2 animals treated with control, *pie-1*, or *top-2* RNAi to probe for *vet-6* mRNA (white). DNA was stained using Hoechst-33342 (blue). P2-associated mRNA signal appears after RNAi depletion of *pie-1* and *top-2*. White dashed circles represent the EMS cell and the red dashed circles represent P2. Scale bar represents a length of 5 µm. (B) HCR was performed using 4-cell embryos treated with *capg-2* RNAi to probe for *vet-6* mRNA (white). DNA was stained using Hoechst-33342 (blue). White dashed circles represent the EMS cell and the red dashed circles represent P2. Scale bar represents a length of 5 µm.

**Figure S3:**
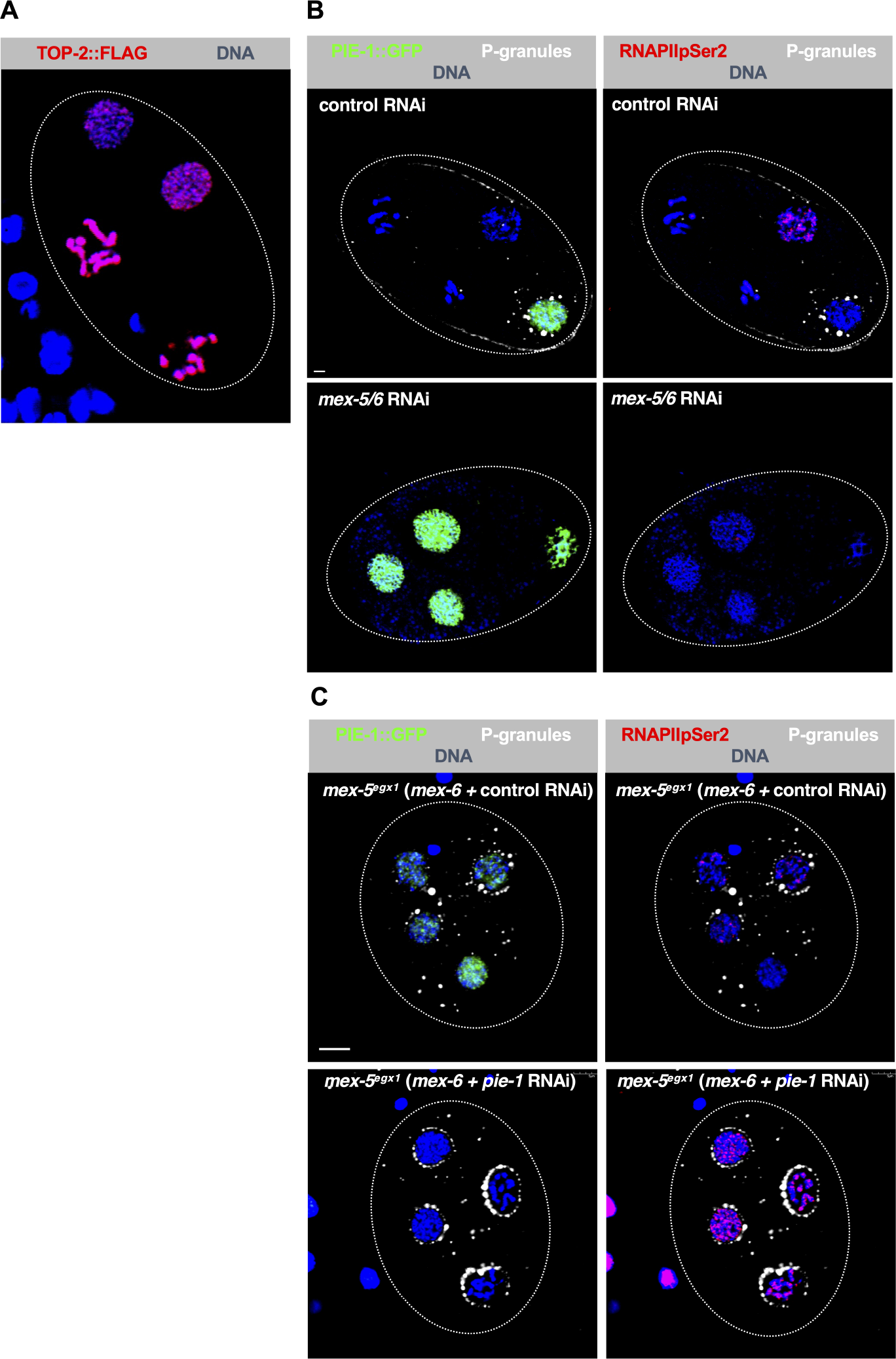
Depletion of MEX-5 and MEX-6 results in PIE-1 expression in all blastomeres of early embryos. (A) Four-cell AG275 embryo fixed and stained for TOP-2::FLAG (red) and DNA (blue). TOP-2 is distributed across all 4 cells of the embryos. Scale bar represents a length of 2 µm. (B) Four-cell embryos from WM330 animals optionally treated with either control or *mex- 5/6* RNAi were fixed and stained for PIE-1::GFP (green), RNAPIIpSer2 (red), DNA (blue), and P-granules (white). Images of the same embryo stained for PIE-1::GFP and RNAPIIpSer2 are shown side by side. Depletion of *mex-5/6* results in the loss of asymmetric distribution of PIE-1 to P2. Scale bar represents a length of 2 µm. (C) Four-cell embryos from EGD175 animals were treated with a combination of either *mex- 6*/control or *mex-6/pie-1* RNAi, then fixed and stained for PIE-1::GFP (green), RNAPIIpSer2 (red), DNA (blue), and P-granules (white). *mex-5* mutant embryos depleted of *mex-6* and *pie-1* produce aberrant RNAPIIpSer2 signal in all blastomeres. Scale bar represents a length of 5 µm.

## References

1. Walker AK, Boag PR, Blackwell TK. Transcription reactivation steps stimulated by oocyte maturation in C. elegans. Dev Biol. 2007 Apr 1;304(1):382–93.

2. Belew MD, Chien E, Michael WM. Characterization of factors that underlie transcriptional silencing in C. elegans oocytes. PLoS Genet. 2023 Jul 21;19(7):e1010831.

3. Chien E, Michael WM. Transcriptional repression during spermatogenesis in C. elegans requires TOP- 2, condensin II, and the MET-2 H3K9 methyltransferase. MicroPubl Biol. 2023 Aug 23;2023:10.17912/micropub.biology.000933.

4. Guven-Ozkan T, Nishi Y, Robertson SM, Lin R. Global transcriptional repression in C. elegans germline precursors by regulated sequestration of TAF-4. Cell. 2008 Oct 3;135(1):149–60.

5. Mello CC, Schubert C, Draper B, Zhang W, Lobel R, Priess JR. The PIE-1 protein and germline specification in C. elegans embryos. Nature. 1996 Aug 22;382(6593):710-2.

6. Seydoux G, Mello CC, Pettitt J, Wood WB, Priess JR, Fire A. Repression of gene expression in the embryonic germ lineage of C. elegans. Nature. 1996 Aug 22;382(6593):713-6.

7. Schaner CE, Deshpande G, Schedl PD, Kelly WG. A conserved chromatin architecture marks and maintains the restricted germ cell lineage in worms and flies. Dev Cell. 2003 Nov;5(5):747–57.

8. Belew MD, Chien E, Wong M, Michael WM. A global chromatin compaction pathway that represses germline gene expression during starvation. J Cell Biol. 2021 Sep 6;220(9):e202009197.

9. Palancade B, Bensaude O. Investigating RNA polymerase II carboxyl-terminal domain (CTD) phosphorylation. Eur J Biochem. 2003 Oct;270(19):3859–70.

10. Shakes DC, Wu JC, Sadler PL, Laprade K, Moore LL, Noritake A, et al. Spermatogenesis-specific features of the meiotic program in Caenorhabditis elegans. PLoS Genet. 2009 Aug;5(8):e1000611.

11. Bowman EA, Bowman CR, Ahn JH, Kelly WG. Phosphorylation of RNA polymerase II is independent of P-TEFb in the C. elegans germline. Development. 2013 Sep;140(17):3703–13.

12. Seydoux G, Dunn MA. Transcriptionally repressed germ cells lack a subpopulation of phosphorylated RNA polymerase II in early embryos of Caenorhabditis elegans and Drosophila melanogaster. Development. 1997 Jun;124(11):2191–201.

13. Tocchini C, Keusch JJ, Miller SB, Finger S, Gut H, Stadler MB, et al. The TRIM-NHL protein LIN-41 controls the onset of developmental plasticity in Caenorhabditis elegans. PLoS Genet. 2014 Aug;10(8):e1004533.

14. Fassnacht C, Tocchini C, Kumari P, Gaidatzis D, Stadler MB, Ciosk R. The CSR-1 endogenous RNAi pathway ensures accurate transcriptional reprogramming during the oocyte-to-embryo transition in Caenorhabditis elegans. PLoS Genet. 2018 Mar;14(3):e1007252.

15. Butuči M, Williams AB, Wong MM, Kramer B, Michael WM. Zygotic Genome Activation Triggers Chromosome Damage and Checkpoint Signaling in C. elegans Primordial Germ Cells. Dev Cell. 2015 Jul 6;34(1):85–95.

16. Wong MM, Belew MD, Kwieraga A, Nhan JD, Michael WM. Programmed DNA Breaks Activate the Germline Genome in Caenorhabditis elegans. Dev Cell. 2018 Aug 6;46(3):302–315.e5.

17. Kim H, Ishidate T, Ghanta KS, Seth M, Conte D, Shirayama M, et al. A co-CRISPR strategy for efficient genome editing in Caenorhabditis elegans. Genetics. 2014 Aug;197(4):1069–80.

18. Gauvin TJ, Han B, Sun MJ, Griffin EE. PIE-1 Translation in the Germline Lineage Contributes to PIE-1 Asymmetry in the Early Caenorhabditis elegans Embryo. G3 (Bethesda). 2018 Dec 10;8(12):3791-801.

19. Choi HM, Calvert CR, Husain N, Huss D, Barsi JC, Deverman BE, et al. Mapping a multiplexed zoo of mRNA expression. Development. 2016 Oct 1;143(19):3632–7.

20. Jaramillo-Lambert A, Fabritius AS, Hansen TJ, Smith HE, Golden A. The Identification of a Novel Mutant Allele of topoisomerase II in Caenorhabditis elegans Reveals a Unique Role in Chromosome Segregation During Spermatogenesis. Genetics. 2016 Dec;204(4):1407–22.

21. Schubert CM, Lin R, de Vries CJ, Plasterk RH, Priess JR. MEX-5 and MEX-6 function to establish soma/germline asymmetry in early C. elegans embryos. Mol Cell. 2000 Apr;5(4):671–82.

22. Chan RC, Severson AF, Meyer BJ. Condensin restructures chromosomes in preparation for meiotic divisions. J Cell Biol. 2004 Nov 22;167(4):613–25.

23. Stanvitch G, Moore LL. cin-4, a gene with homology to topoisomerase II, is required for centromere resolution by cohesin removal from sister kinetochores during mitosis. Genetics. 2008 Jan;178(1):83–97.

24. Bembenek JN, Verbrugghe KJ, Khanikar J, Csankovszki G, Chan RC. Condensin and the spindle midzone prevent cytokinesis failure induced by chromatin bridges in C. elegans embryos. Curr Biol. 2013 Jun 3;23(11):937–46.

25. Batchelder C, Dunn MA, Choy B, Suh Y, Cassie C, Shim EY, et al. Transcriptional repression by the Caenorhabditis elegans germ-line protein PIE-1. Genes Dev. 1999 Jan 15;13(2):202–12.

26. Wang JT, Seydoux G. Germ cell specification. Adv Exp Med Biol. 2013;757:17–39.

27. Kim H, Ding YH, Lu S, Zuo MQ, Tan W, Conte D Jr, Dong MQ, Mello CC. PIE-1 SUMOylation promotes germline fates and piRNA-dependent silencing in C. elegans. Elife. 2021 May 18;10:e63300.

