## Supplementary material for "The condensin II/TOP-2 axis silences transcription during germline specification in *C. elegans*": BioRxiv Supporting.docx

**Supporting information figures and legends**

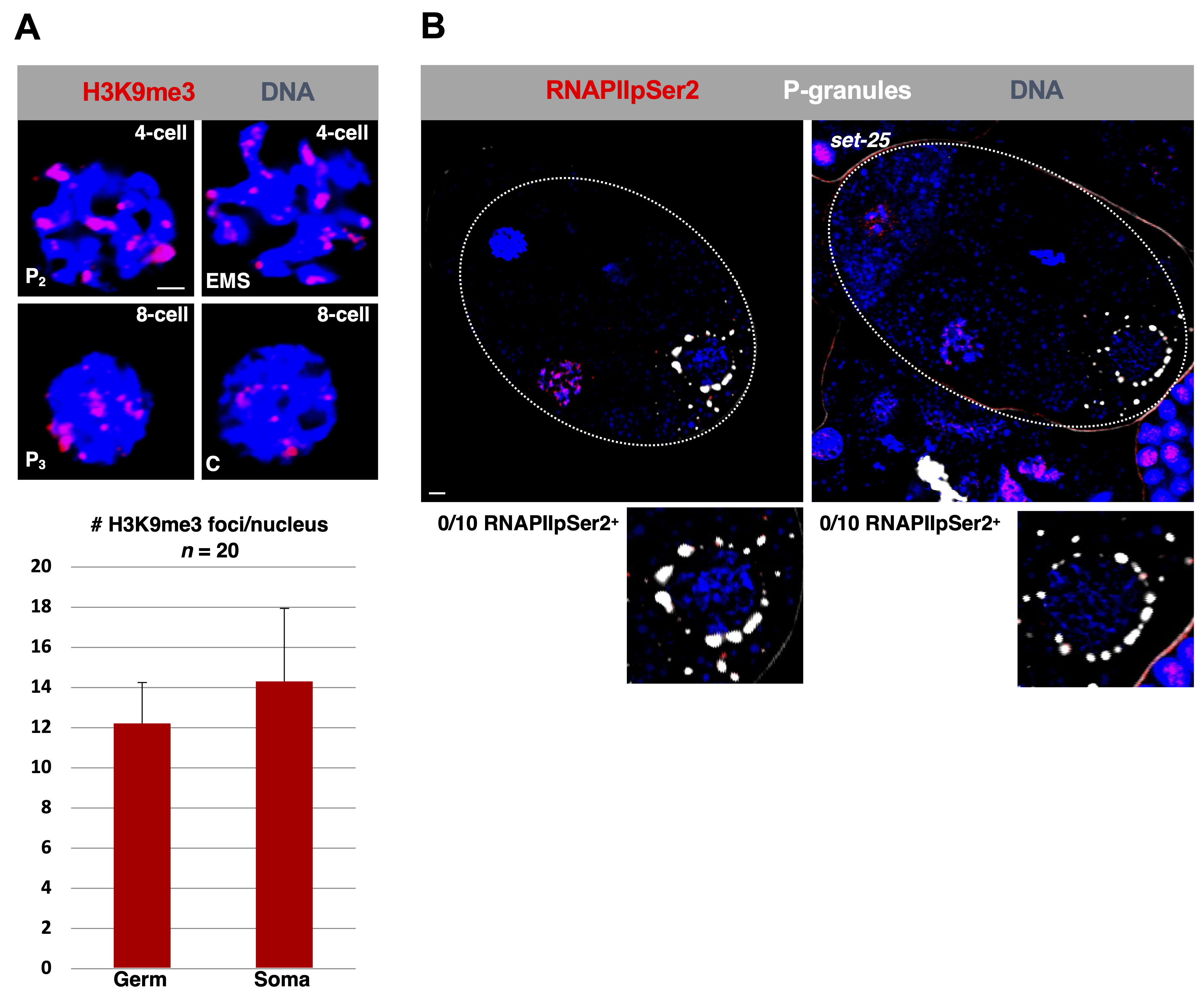

**Figure S1: RNAPII-mediated transcription is still repressed in P_2_ after loss of SET-25.**

(A) Four- and eight-cell embryos were examined were fixed and stained for H3K9me3 (red) and DNA (blue). Quantification of H3K9me3 foci for P_2_ and P_3_ (Germ) or EMS and C (Soma) is shown below. Scale bar represents a length of 2 µm.
(B) Four-cell embryos, either wild type (N2) or *set-25* mutants (strain MT17463), were fixed and stained for RNAPIIpSer2 (red) and P-granules (blue). 10 samples were analyzed. Both wild-type and *set-25* mutants lacked RNAPIIpSer2 signal in P_2_. Scale bar represents a length of 2 µm.

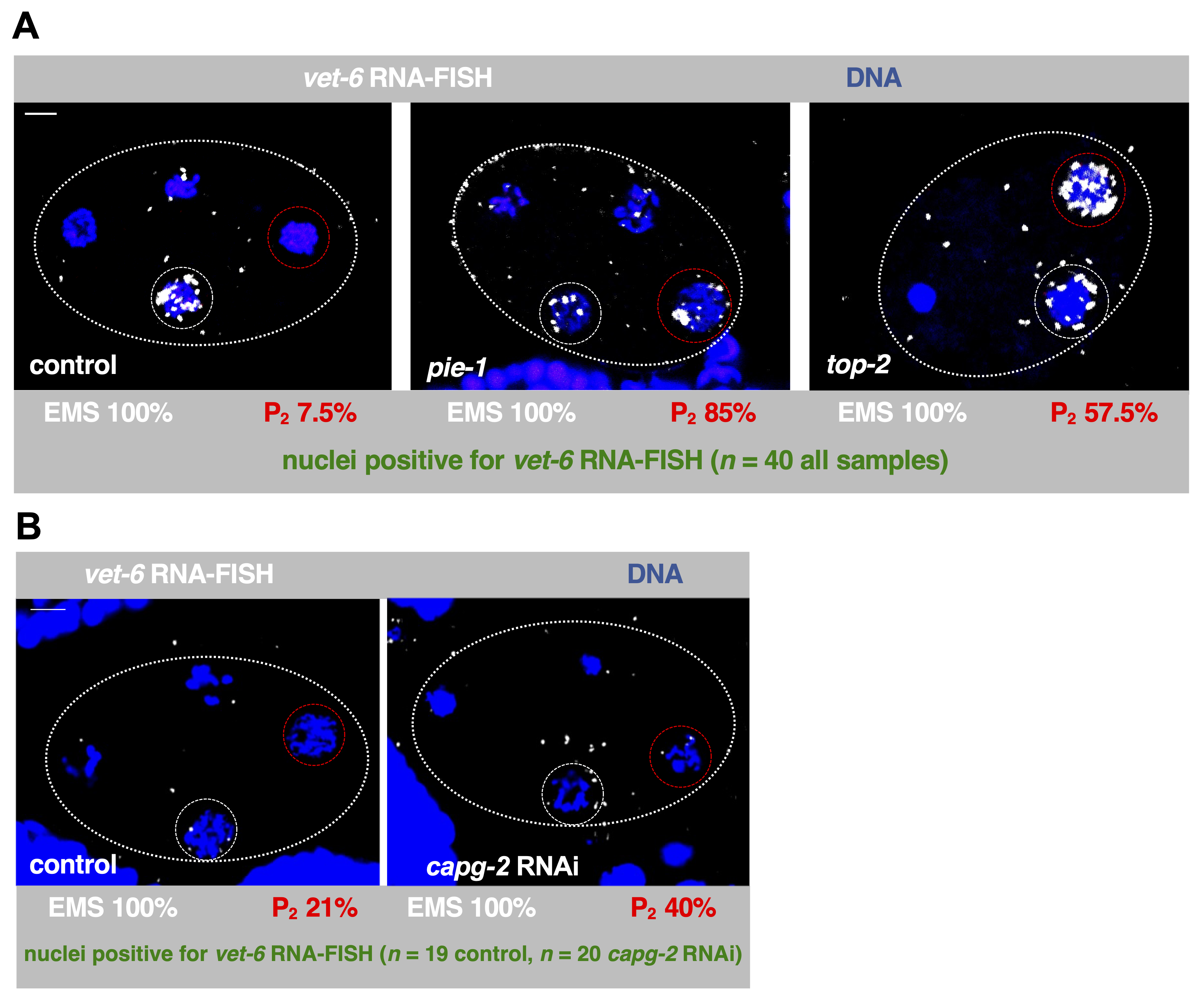

**Figure S2: EMS specific gene is aberrantly expressed in the P_2_ cell of 4-cell embryos after loss of TOP-2 but not CAPG-2.**

(A) HCR was performed using 4-cell embryos from N2 animals treated with control, *pie-1*, or *top-2* RNAi to probe for *vet-6* mRNA (white). DNA was stained using Hoechst-33342 (blue). P_2_-associated mRNA signal appears after RNAi depletion of *pie-1* and *top-2*. White dashed circles represent the EMS cell and the red dashed circles represent P_2_. Scale bar represents a length of 5 µm.
(B) HCR was performed using 4-cell embryos treated with *capg-2* RNAi to probe for *vet-6* mRNA (white). DNA was stained using Hoechst-33342 (blue). White dashed circles represent the EMS cell and the red dashed circles represent P_2_. Scale bar represents a length of 5 µm.

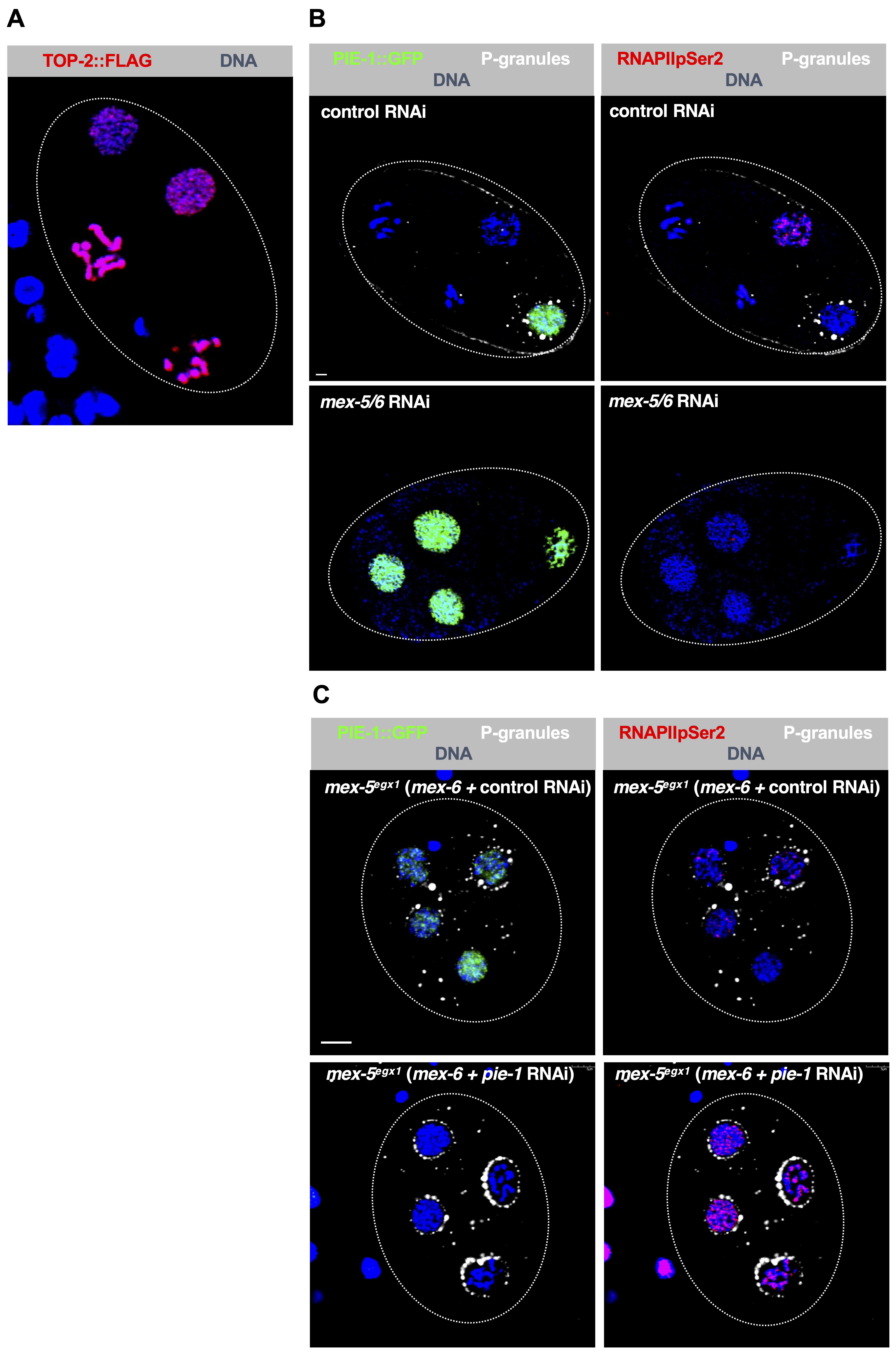

**Figure S3: Depletion of MEX-5 and MEX-6 results in PIE-1 expression in all blastomeres of early embryos.**

(A) Four-cell AG275 embryo fixed and stained for TOP-2::FLAG (red) and DNA (blue). TOP-2 is distributed across all 4 cells of the embryos. Scale bar represents a length of 2 µm.
(B) Four-cell embryos from WM330 animals optionally treated with either control or *mex-5/6* RNAi were fixed and stained for PIE-1::GFP (green), RNAPIIpSer2 (red), DNA (blue), and P-granules (white). Images of the same embryo stained for PIE-1::GFP and RNAPIIpSer2 are shown side by side. Depletion of *mex-5/6* results in the loss of asymmetric distribution of PIE-1 to P_2_. Scale bar represents a length of 2 µm.
(C) Four-cell embryos from EGD175 animals were treated with a combination of either *mex-6*/control or *mex-6/pie-1* RNAi, then fixed and stained for PIE-1::GFP (green), RNAPIIpSer2 (red), DNA (blue), and P-granules (white). *mex-5* mutant embryos depleted of *mex-6* and *pie-1* produce aberrant RNAPIIpSer2 signal in all blastomeres. Scale bar represents a length of 5 µm.
